# A mass-transfer-constrained thermodynamic model reveals how fluid flow reshapes interspecies electron transfer during syntrophic propionate oxidation

**DOI:** 10.64898/2026.06.25.734252

**Authors:** Yuan Fang, Ran Mei

**Author notes:** Corresponding author: Ran Mei, Department of Civil and Environmental Engineering, University of Illinois Urbana-Champaign, 205 North Mathews Ave, Urbana, Illinois 61801, USA.

## Abstract

Syntrophic propionate oxidation in methanogenic environments depends on interspecies electron transfer via hydrogen and formate, yet the physical factors governing the relative use of these electron carriers remain poorly understood. Here, we investigated how fluid flow reshapes electron-transfer energetics and pathway expression in the obligately syntrophic propionate oxidizer *Pelotomaculum schinkii* grown in coculture with *Methanospirillum hungatei*. We combined coculture physiology, transcriptomics, and a mass-transfer-constrained thermodynamic model to estimate H₂ and formate concentrations near the *P. schinkii* cell surface and calculate the Gibbs free-energy changes associated with H₂- and formate-mediated propionate oxidation under different mixing conditions. Under the unmixed condition, formate-mediated propionate oxidation was thermodynamically more favourable than the H₂-mediated pathway, consistent with elevated expression of genes involved in formate production. Mixing altered both process energetics and gene expression. H₂-mediated propionate oxidation was more sensitive to mixing than the formate-mediated pathway. The expression of major hydrogenases and formate dehydrogenases closely tracked these pathway-specific energetic shifts. The relative use of formate- and H₂-mediated electron transfer by syntrophs also varied in anaerobic digesters. Together, these findings provide new insights into syntrophic propionate degradation and its thermodynamic constraints.

## Introduction

Anaerobic microbial communities are widespread and contribute fundamentally to global carbon cycling and methane production ^1^. Complete methanogenic degradation of many organic compounds requires syntrophy, in which reducing equivalents generated during substrate oxidation are transferred to a methanogenic partner by shuttle components ^2^. The shuttle components must be maintained at low concentrations to relieve the thermodynamic constraints on substrate oxidation ^3^. Molecular hydrogen and formate are the principal electron shuttle components for syntrophy ^4–6^. They are often considered interchangeable ^3,7^ because they share similar redox potentials under standard and biologically relevant conditions, and have been widely demonstrated to be simultaneously involved in syntrophy ^6,8–11^.

Increasing evidence indicates that the relative use of H₂ and formate in syntrophy is physiologically regulated rather than random or fixed. When *Syntrophomonas wolfei* grows on butyrate in coculture with *Methanospirillum hungatei*, one proteomic study found formate dehydrogenases to be predominant ^12^, whereas other studies suggested that certain culture conditions may favor the use of hydrogenase ^13,14^. Similarly, transcriptomic and proteomic analyses of *Syntrophobacterium fumaroxidans* (previously known as *Syntrophobacter fumaroxidans* ^15^) suggest that the relative use of hydrogenases and formate dehydrogenases varies with substrate and growth mode ^16,17^. Such flexibility has also been observed in *Desulfovibrio alaskensis* G20 during syntrophic growth on lactate with methanogens under sulfate-free conditions, where the relative contributions of H₂ and formate to interspecies electron transfer varied depending on the methanogenic partner and dilution rate ^18^. Together, these findings indicate that syntrophs can flexibly shift between H₂ and formate as electron carriers, although the environmental factors governing this shift remain poorly understood.

One potential environmental factor is fluid flow which may differentially affect H_2_ and formate transport. In the aqueous phase, H₂ diffuses approximately three times faster than formate, whereas its solubility is about two orders of magnitude lower ^19–21^. At the gas-liquid interface, H₂ partitions between the two phases, providing an additional transport pathway that is not applicable to ionic formate at near-neutral pH. At the cellular level, H₂ readily permeates cell membranes, whereas formate generally requires dedicated transport proteins for transmembrane movement. These physicochemical differences subject H_2_ and formate to distinct transport constraints and may cause them to respond differently to fluid flow.

Fluid flow can modify these transport constraints by altering solute transport between cells and the surrounding environment. Increased fluid velocity enhances mass transfer to and from suspended microorganisms ^22,23^. In syntrophic communities, fluid flow may therefore influence the removal of H₂ and formate from producing cells, their delivery to consumer cells, their redistribution within the bulk liquid, and, for H₂, exchange between the liquid and gas phases. In addition, fluid flow can affect cell proximity, aggregation, and spatial organization, all of which are known to influence syntrophic metabolism ^24–26^.

We hypothesized that fluid flow influences the relative use of H₂- and formate-mediated electron transfer during syntrophic degradation of propionate. To test this hypothesis, we used a coculture of *Pelotomaculum schinkii* HH and *Methanospirillum hungatei* JF-1 that degrades propionate syntrophically. Propionate was chosen because it is a major intermediate in anaerobic organic matter decomposition, and its oxidation is among the most thermodynamically challenging reactions in anaerobic metabolism ^27^. Both H₂- and formate-mediated propionate oxidation are highly endergonic under standard conditions:

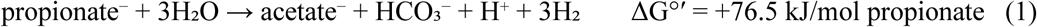

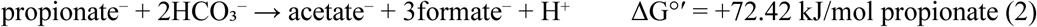

These severe thermodynamic constraints make syntrophic propionate oxidation particularly sensitive to the efficiency of interspecies electron transfer ^6,27,28^. *P. schinkii* was selected because it is an obligately syntrophic propionate oxidizer ^29^ whose genome encodes multiple hydrogenases and formate dehydrogenases ^30^, suggesting the capacity to flexibly use H₂ and/or formate as the electron carrier. To determine whether fluid flow alters the relative use of H₂ and formate, we evaluated cocultures under different mixing conditions by integrating physicochemical measurements, comparative genomic analysis, transcriptomics, and a novel mass-transfer-constrained thermodynamic model. Importantly, the mass-transfer-constrained model enabled estimation of H₂ and formate concentrations in the immediate vicinity of syntroph cells, potentially improving estimates of local thermodynamic conditions compared with those based on bulk-liquid concentrations ^31,32^.

## Materials and Methods

### Cultivation experiment

The coculture of *P. schinkii* HH and *M. hungatei* JF-1 (DSM 15200) was obtained from DSMZ (Braunschweig, Germany). Cultures were grown anaerobically in serum bottles (160 mL) with modified Widdel medium prepared following established protocols ^33^. Propionate was provided at a final concentration of 20 mM. Anaerobic techniques were adapted from previously described methods ^34^. Cultures were incubated at 37 °C. Mixing conditions were controlled by magnetic stirring at 0, 130, or 250 rpm using cylindrical stir bars (30 × 6 mm) and a magnetic stirring system. Triplicate bottles were prepared for each condition. Detailed methods are provided in the Supplementary Information.

### GC measurement

Headspace gas composition was measured using a GC-2030 gas chromatograph (Shimadzu, Kyoto, Japan) equipped with a thermal conductivity detector (TCD), and a flame ionization detector (FID). H₂ concentrations below the detection limit of the GC-TCD method (50 ppm, approximately 5 Pa) were treated as nondetects and assigned the detection-limit value for subsequent calculations ^35^. Propionate and acetate in the liquid phase were quantified by GC-FID after acidification and methyl tert-butyl ether (MTBE) extraction with iso-butyric acid used as an internal standard, adapted from a previous study ^36^. Because formate concentrations were near or below the detection limit of the GC-FID method, selected samples were analysed by GC-MS at the Roy J. Carver Biotechnology Center at the University of Illinois.

### RNA sequencing

Biomass from cocultures was harvested at two time points: day 2 (∼2 mM propionate consumed at 0 and 130 rpm, with less than 0.5 mM consumption at 250 rpm) and day 8 (∼10 mM consumed at 0 and 130 rpm and ∼5 mM at 250 rpm) with biological triplicates. Cultures were harvested anaerobically by centrifugation, stabilised with RNAprotect Bacteria Reagent (QIAGEN, Hilden, Germany), and stored at −80 °C. RNA was extracted using the QIAGEN RNeasy Mini Kit with bead-beating lysis and sequential DNase treatments. Ribosomal RNA depletion, strand-specific library preparation, and sequencing on an Illumina NovaSeq X Plus platform using 2 × 151-bp paired-end reads were performed at the Roy J. Carver Biotechnology Center.

### Transcriptomic analysis

Reference genomes of *P. schinkii* HH (GCF_004369205.1) and *M. hungatei* JF-1 (GCF_000013445.1) were downloaded from NCBI and annotated using Prokka 1.15.6 ^37^. Genes encoding hydrogenases and formate dehydrogenases were subsequently identified and manually curated based on protein-domain annotations, sequence similarity to characterised proteins, subunit composition, and genomic context. RNA-seq reads were quality filtered using BBTools v39.52 ^38^, mapped to the two reference genomes using BWA-MEM2 v2.2.1 ^39^, and quantified at the coding-sequence level using featureCounts v2.1.1 ^40^ with strand-specific counting (–s 2). Gene expression was calculated separately for each organism as transcripts per million (TPM) and normalised to the organism-specific median gene expression, following a previous study ^41^. Gene ranks were determined from the mean normalised expression across biological replicates.

### Microscope

For cell observation, samples collected from cocultures on days 2 and 8 were fixed in 4% paraformaldehyde at 4 °C for 4 h. Fixed cells were washed twice with 1× PBS and stored in 50% ethanol prepared in 1× PBS at −20 °C until imaging. Before imaging, cells were washed with 1× PBS to remove ethanol and mounted with DAPI-Fluoromount-G™ (Electron Microscopy Sciences, Hatfield, PA, USA) according to the manufacturer’s instructions. Mounted samples were air-dried for 5 min and imaged using an Axio Observer.Z1 epifluorescence microscope (Carl Zeiss, Oberkochen, Germany).

### Quantitative PCR

*P. schinkii* cell density in cocultures was quantified using qPCR. DNA was extracted from coculture samples (50 mL) collected at day 2 and day 8 using the QIAGEN DNeasy PowerLyzer Microbial Kit according to the manufacturer’s instructions. A previously developed primer set ^42^ targeting *Firmicutes* 16S rRNA gene (934F, 5′-GGAGYATGTGGTTTAATTCGAAGCA-3′; 1040R, 5′-ACCATGCACCACCTGTC-3′) was used. Biological triplicates were analysed for each condition. qPCR was performed on a StepOnePlus Real-Time PCR System (Applied Biosystems, Waltham, MA, USA) using Luna Universal qPCR Master Mix (New England Biolabs, Ipswich, MA, USA) in 20 µL reaction volumes. Amplification consisted of 95 °C for 20 s, followed by 40 cycles of 95 °C for 3 s and 61.5 °C for 30 s. Standard curve was generated from serial dilutions of a synthetic double-stranded DNA standard containing the target amplicon region (Integrated DNA Technologies, Coralville, IA, USA). Cell densities were calculated by considering four 16S rRNA gene copies per *P. schinkii* genome.

### Inhibition experiment

To estimate the relative electron flow through H₂ and formate during syntrophic propionate oxidation, we applied an inhibition-based approach reported previously ^18^. Methanogenesis was inhibited with 2-bromoethanesulfonate (BES) to suppress H₂ and formate consumption by *M. hungatei*, allowing H₂ and formate produced by *P. schinkii* to accumulate for quantification. Cocultures were cultivated under the same conditions as described above. BES was added to a final concentration of 10 mM ^43^. Following BES addition, headspace H₂ and dissolved formate concentrations were measured after 24 h ^44^. The electron partitioning between H₂ and formate was calculated as Δn_H2_ /(Δn_H2_ + Δn_formate_) and Δn_formate_/(Δn_H2_ + Δn_formate_), respectively, where Δn represents the increase in amount following BES addition. These estimates assumed negligible consumption and interconversion of H₂ and formate following inhibition. Total H₂ included both headspace and dissolved fractions, with the latter estimated using Henry’s law.

### Mass-transfer-constrained thermodynamic modelling

Based on the classical liquid-film mass-transfer framework ^45,46^, a mass-transfer-constrained thermodynamic model was used to estimate H₂ and formate concentrations at the surface of *P. schinkii* cells and free energy associated with H₂ and formate production (Table 1, detailed calculation in Supplementary Information). For each electron carrier *i* (H₂ or formate), the surface-production flux was balanced by external mass transfer assuming steady state: *J_i_* = *k_L_*_,*i*_(*C_surface_*_,*i*_ − *C_bulk_*_,*i*_), where *J_i_* is the cell-surface production flux, *k_L_*_,*i*_ is the liquid-side carrier-specific mass transfer coefficient, and *C_surface_*_,*i*_ and *C_bulk_*_,*i*_ are the cell-surface and bulk-liquid concentrations, respectively. This allows the estimation of cell-surface concentration as: 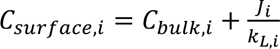. Bulk concentrations were determined at the corresponding sampling times and mixing conditions. Formate was measured directly, whereas dissolved H₂ was calculated from headspace partial pressure using Henry’s law ^47^. *J_i_* was estimated from interval-specific propionate consumption, the stoichiometric production of electron carrier equivalents per mole of propionate, the fraction of electrons allocated to catabolism ^48,49^, the electron partitioning between H_2_ and formate determined in inhibition experiment, qPCR-derived *P. schinkii* cell density, and cell surface area estimated from the electron microscopic image from a previous study ^30^. *k_L_*_,*i*_ was calculated from molecular diffusion coefficient, volume-equivalent cell radius, and Sherwood number ^22^. Sherwood number was calculated from Ranz–Marshall correlation ^50,51^ with Reynolds number and Schmidt number using the stir-bar tip velocity as a characteristic fluid velocity. All the data used in model calculation are included in Supplementary Information.

**Table 1.** Summary of parameters used in the mass-transfer-constrained thermodynamic analysis.

| Category | Parameter | Symbol/unit | Determination or source |
|---|---|---|---|
| Measured in this study | Headspace CH<sub>4</sub> and H<sub>2</sub> | Pa | GC-TCD |
|  | Bulk liquid propionate and acetate | mM | GC-FID |
|  | Bulk liquid formate | μM | GC-MS |
|  | Cell abundance | cell/mL | qPCR |
|  | Electron partitioning between H<sub>2</sub> and formate | dimensionless | Inhibition-based partitioning<sup>18</sup> |
| Calculated in this study | Dissolved bulk H<sub>2</sub> concentration | μM | Calculated from headspace partial pressure using Henry's law<sup>47</sup> |
|  | Fraction of electrons allocated to catabolism | fe, dimensionless | Calculated from thermodynamic electron-equivalent approach<sup>49</sup> |
|  | Sherwood number | Sh, dimensionless | Calculated using the Ranz–Marshall correlation<sup>50,51</sup> |
|  | Liquid-side mass-transfer coefficient | kL, m/s | Calculated from Sherwood number definition<sup>22</sup> |
| Literature constants | Henry's law constants | L•atm/mol | Reference<sup>47</sup> |
|  | Cell dimensions | μm | Reference<sup>30</sup> |
|  | Standard redox potentials | E<sub>0</sub>', V | Reference<sup>53</sup> |
|  | Standard free energies of formation | ΔG<sub>f</sub><sup>0</sup>, kJ/mol | Reference<sup>52</sup> |
|  | Half-reaction free energies | ΔG<sub>0</sub>', kJ/mol | Reference<sup>48</sup> |

Estimated surface concentrations were then used to calculate the Δ*G* of H₂- and formate-mediated propionate oxidation using Δ*G* = Δ*G°’* + *RT*ln *Q*, where Δ*G°’* is calculated from standard Gibbs energies of formation and standard redox potentials ^52,53^; R is the ideal gas constant, 8.314 J/(mol·K); T is the incubation temperature, 310 K; and Q is the reaction quotient calculated using the estimated cell-surface H₂ or formate concentration determined above.

Model robustness was evaluated by independently varying uncertain transport and biological parameters. The liquid-side mass-transfer coefficient was tested under both the baseline advection-plus-diffusion formulation and a diffusion-only scenario (Sherwood number=2). Cell surface area, assigned H₂ concentrations below the detection limit, substrate-consumption rate, and electron partitioning between H_2_ and formate were varied by ±25%, while the electron fraction allocated to catabolism was varied from 0.86 to 1.00. Pathway-specific Gibbs free-energy changes were recalculated for all mixing conditions and sampling times and were compared with the baseline scenario.

### Full-scale digester metatranscriptomic analysis

Published time-series metatranscriptomic datasets from full-scale anaerobic digesters ^54,55^ were analysed to assess the relative expression of formate dehydrogenases and hydrogenases in real-world methanogenic environments. These digesters were continuously operated, suspended-growth systems, and most were operated under mesophilic conditions with sludge retention times of 20–30 days. Multiple metatranscriptomic samples were obtained by the original authors from each digester at different time points. The dataset from Kieft et al. ^54^ included 14 metatranscriptomes from one digester, while the dataset from Nobu et al. ^55^ included three metatranscriptomes from each of nine digesters. Detailed operating conditions, sampling information, and data accession numbers are provided in the Supplementary Information. Metatranscriptomic reads were mapped to four representative syntrophic reference genomes: *P. schinkii* (GCF_004369205.1), *Syntrophobacterium fumaroxidans* (GCF_000014965.1), *Syntrophus aciditrophicus* (GCF_000013405.1), and *Syntrophomonas wolfei* (GCF_000014725.1). Because the metatranscriptomic libraries were unstranded, read counting was performed in unstranded mode (-s 0). Transcript abundances were expressed as TPM. For each reference syntroph in each sample, the formate dehydrogenase:hydrogenase expression ratio was calculated as the summed TPM of all the formate dehydrogenase genes in the genome divided by the summed TPM of all the hydrogenase genes. For the data from Kieft et al. ^54^ where more physicochemical and operational data are available, redundancy analysis was performed in R using the vegan package ^56^ to examine associations between syntroph-specific formate dehydrogenase: hydrogenase expression ratios and AD process parameters. Selected process parameters, including nitrite, hydraulic retention time, nitrate, ammonia, soluble chemical oxygen demand, volatile fatty acids, temperature, pH, biogas CH_4_ and CO_2_ percentages were standardised before analysis. Model significance was assessed using 999 permutations.

## Results

### Electron-transfer capacity of P. schinkii HH

The genome of *P. schinkii HH* was analysed to understand its electron transfer capacity. Consistent with a previous report ^30^, *P. schinkii* HH oxidizes propionate through the methylmalonyl-CoA pathway and disposes of electrons through H₂ and/or formate. Its genome encodes multiple hydrogenases and formate dehydrogenases (Figure 1). These include an electron-confurcating [FeFe]-hydrogenase Hyd, an energy-conserving [NiFe]-hydrogenase Ech, an electron-confurcating formate dehydrogenase Fdh-Hyl, and a monomeric formate dehydrogenase FdhH. There are several auxiliary electron transduction proteins, including Rnf, Fix, and Nfn, which may redistribute electrons among ferredoxin, NAD(H), NADP(H), and the quinone pool.

**Figure 1.**
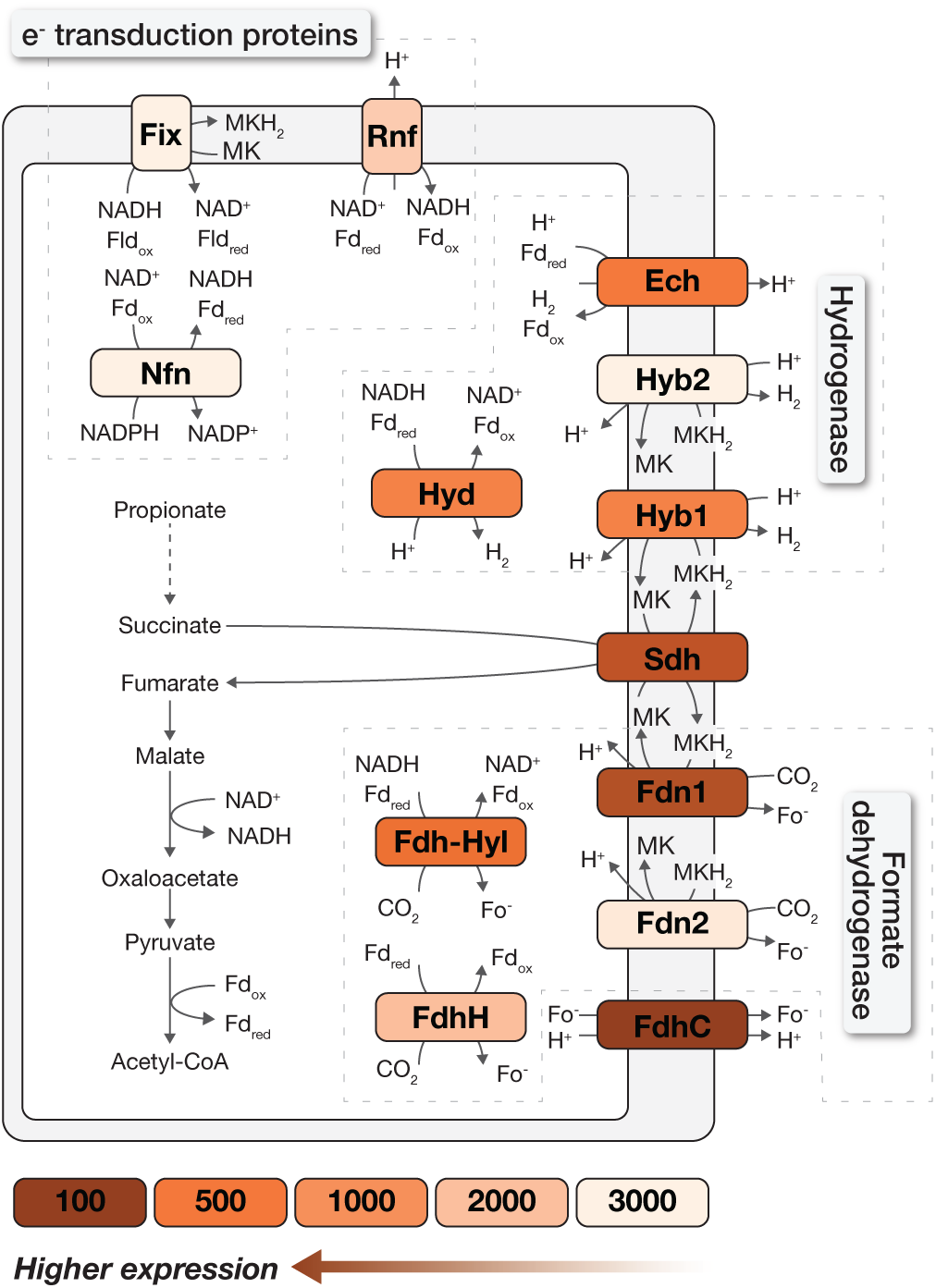
Predicted electron-transfer capacity of *P. schinkii* HH. Box colors indicate the average rank of all subunit genes of each complex under unmixed condition at day 2. Numbers in the color key are rank values; darker colors and lower ranks indicate higher transcript abundance. Schematic of predicted electron-transfer pathways in *P. schinkii* HH, showing enzymes involved in hydrogen and formate production and their connections to propionate oxidation. Enzyme complexes are color-coded by their average gene expression ranks under unmixed conditions at day 2, with darker colors indicating higher transcript abundance.

In addition to these well-known electron transfer proteins, *P. schinkii* HH harbors a distinctive locus that encodes three membrane-associated menaquinol-linked hydrogenases and formate dehydrogenases (Psch_RS11065–11145, Figure S1). The locus encodes an HybABCO-type [NiFe]-hydrogenase, designated Hyb1, together with two heterotrimeric formate dehydrogenase complexes, Fdn1 and Fdn2. Sequence analysis indicated that the formate-producing catalytic subunits of Fdn1 and Fdn2 are similar to the canonical FdnG, whereas their electron-transfer subunits are more closely related to HybA and HybB than to FdnH and FdnI in *E. coli* ^57^ (Figure S1). This is bioenergetically important because HybB-family proteins can use proton motive force to drive the unfavourable reoxidation of menaquinol through H₂ or formate production ^58,59^, consistent with a classic reverse-electron-transfer mechanism ^3^. Comparative genomic analysis further showed that the related Fdn–Hyb arrangement was conserved in the propionate-degrading *Pelotomaculum* species but was absent from the non-propionate-degrading *Pelotomaculum* species and from closely related *Desulfotomaculum* species capable of non-syntrophic propionate degradation.

### Mass-transfer-constrained thermodynamics explains the preferential use of formate under the unmixed condition

To determine how mixing affected syntrophic metabolism and interspecies electron transfer, cocultures were incubated without mixing or with mixing at 130 and 250 rpm. Propionate consumption, acetate formation, and methane production followed the expected stoichiometry of syntrophic propionate oxidation (Figure S2). The apparent propionate consumption rate was similar under the unmixed condition and 130 rpm: 1.02 mmol/(L·day) at 0 rpm and 1.09 mmol/(L·day) at 130 rpm, but decreased to 0.50 mmol/(L·day) at 250 rpm. H₂ and formate concentrations were highest on day 2 and day 4, subsequently decreased, and remained relatively stable, suggesting an apparent steady state between their production and consumption. Microscopic examination showed that no aggregates formed under any tested conditions and time points (Figure S3). Samples were collected on days 2 and 8 under all three mixing conditions. However, these matched sampling times may correspond to different stages of culture development based on differences in propionate degradation: at day 2, approximately 2 mM propionate had been consumed at 0 and 130 rpm, compared with less than 0.5 mM at 250 rpm. At day 8, approximately 10 mM propionate had been consumed at 0 and 130 rpm, compared with approximately 5 mM at 250 rpm.

Under the unmixed condition, the expression profile of *P. schinkii* indicated preferential use of formate-mediated electron transfer at both day 2 and day 8, based on the gene expression ranks (Table S1). At day 2, the menaquinol-linked formate dehydrogenase Fdn1 was among the most highly expressed electron-transfer complexes with a mean rank of 224, whereas the alternative menaquinol-reoxidizing complexes Hyb1, Hyb2, Fdn2, and Fix ranked substantially lower (724-3641; Figure 1). The ferredoxin- and NADH-linked Fdh–Hyl formate dehydrogenase was also more highly expressed than the hydrogenase counterpart Hyd (mean rank 448 and 723, respectively). In addition, the formate transporter FdhC ranked 58th among all genes. Together, these patterns support formate as the dominant electron disposal route under the unmixed condition.

To test whether the transcriptional preference reflected local energetics, we applied a thin-film mass-transfer model to estimate H₂ and formate concentrations near the *P. schinkii* cell surface (Figure 2; brief description of parameters in Table 1; detailed values used for calculation in Table S2-S5). The model assumes that, at steady state, production of an electron carrier is balanced by transport away from the cell:

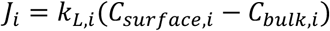

where *J_i_* is the production flux of carrier *i* (H₂ or formate), *k_L_*_,*i*_ is its liquid-side mass-transfer coefficient, and *C_surface_*_,*i*_ and *C_bulk_*_,*i*_ are the near-surface and bulk concentrations, respectively. Under the unmixed condition, the estimated H₂ partial pressure and formate concentration at the cell surface were 16.30 Pa and 12.13 µM, respectively. The corresponding ΔG values were −1.58 kJ/mol propionate for the H₂-producing pathway (reaction 1) and −6.00 kJ/mol propionate for the formate-producing pathway (reaction 2), giving formate an energetic advantage of 4.42 kJ/mol propionate. By comparison, calculations based on the measured bulk concentrations (12.76 Pa for H₂ and 11.86 µM for formate) yielded ΔG values of −3.48 and −6.18 kJ/mol propionate for the H₂- and formate-producing pathways, respectively. Consequently, the energetic advantage of formate production was only 2.70 kJ/mol, indicating that calculations based on bulk concentrations underestimate the thermodynamic advantage of formate-mediated electron transfer.

**Figure 2.**
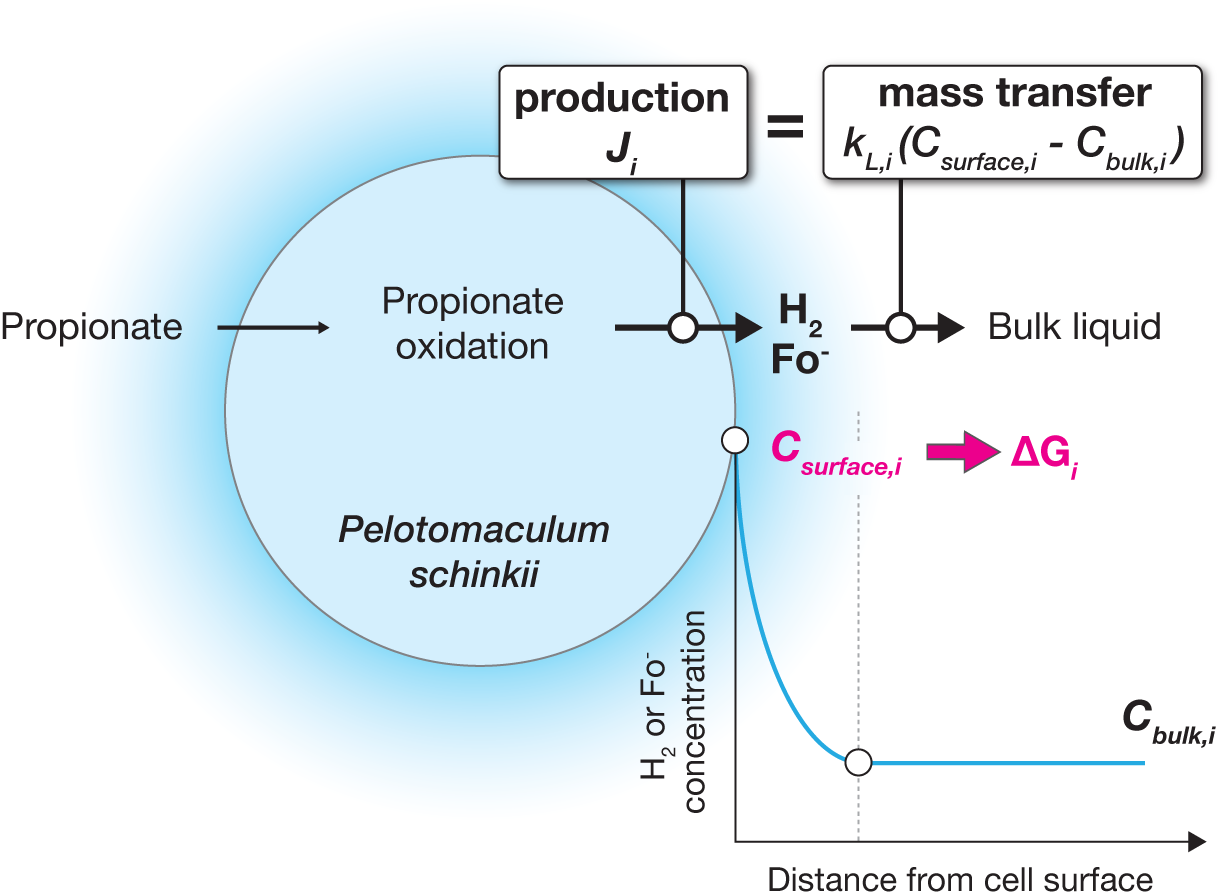
Conceptual representation of the mass-transfer constrained thermodynamic model. Propionate oxidation by *P. schinkii* generates H_2_ and/or formate. At steady state, the carrier-production flux, *J_i_*, is balanced by external mass transfer to the bulk liquid, *k_L,i_* (*C_surface,i_* - *C_bulk,i_*), where *i* denotes H_2_ or formate. The resulting surface concentration is used to calculate ΔG for H_2_ or formate production from propionate. Diagram showing hydrogen and formate produced during propionate oxidation by *P. schinkii* and their transfer into the bulk liquid. A line graph illustrates decreasing concentrations with distance from the cell surface, with surface concentrations used to calculate Gibbs free energy changes (ΔG).

### Mixing shifted the thermodynamic favourability of H₂- and formate-mediated electron transfer

We next asked whether fluid flow altered this favourability of formate. At day 2, mixing at 130 rpm shifted the cell-surface ΔG of the H₂-mediated pathway by 2.02 kJ/mol toward greater favourability relative to the unmixed condition, whereas the formate-mediated pathway became slightly less favourable by 0.21 kJ/mol (Figure 3A; Table S2). This energetic shift coincided with increased expression of the major hydrogenases Hyd and Hyb1 (1.6- to 5.2-fold and 1.2- to 4.1-fold increases, respectively; Table S6), modest changes in Ech expression (0.8- to 1.4-fold), and a general decrease in formate dehydrogenase expression (0.1- to 1.1-fold). Nevertheless, formate-mediated propionate oxidation remained thermodynamically more favourable than the H₂-mediated pathway (−5.80 vs. −3.61 kJ/mol, respectively), with a smaller advantage (2.19 kJ/mol) compared to the unmixed condition (4.42 kJ/mol). A similar pattern was observed at day 8 (Figure 3B; Table S2), when the H₂-mediated pathway exhibited a larger thermodynamic shift (2.74 kJ/mol) than the formate-mediated pathway (0.11 kJ/mol). Consequently, the H₂-mediated pathway became thermodynamically more favourable than the formate-mediated pathway (−3.77 vs. −2.75 kJ/mol, respectively). Consistent with this shift, most hydrogenases were upregulated, except for Ech, whereas the expression of formate dehydrogenases changed little or decreased.

**Figure 3.**
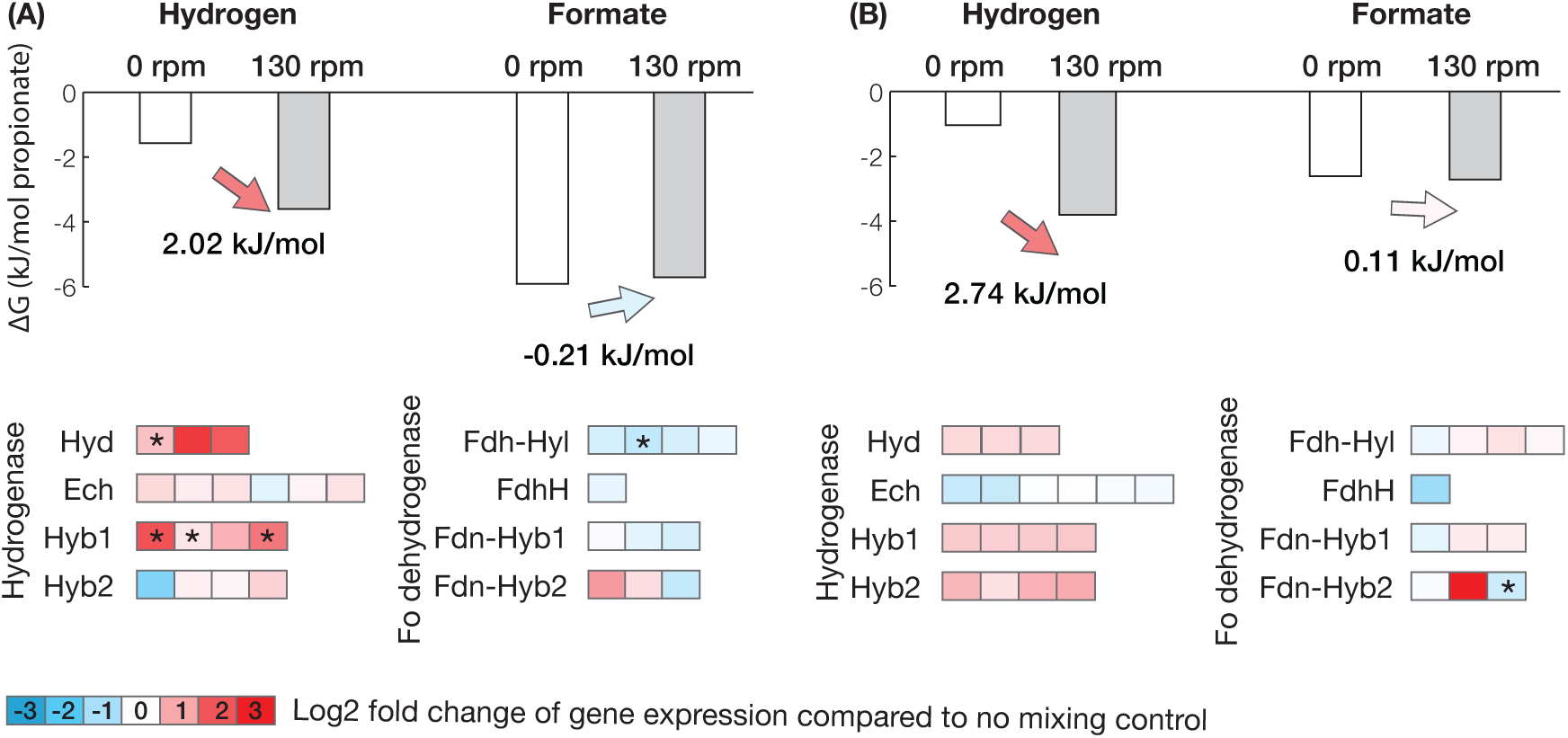
Mixing-dependent changes in the thermodynamic favourability and transcription of H₂- and formate-mediated electron-transfer pathways in *P.* schinkii at 130 rpm. Bars show the calculated Gibbs free-energy change of propionate oxidation through H₂- and formate-mediated pathways under unmixed condition(white) and mixed at 130 rpm (light grey). Panels compare day 2 (A) versus day 8 (B). Arrows indicate the change in ΔG calculated as ΔG_unmixed_ - ΔG_mixed_; positive values indicate that mixing made ΔG more negative and thus more favourable, whereas negative values indicate reduced favourability. Heatmaps below each comparison show the log2 fold change in expression of major hydrogenase and formate dehydrogenase complexes at 130 rpm relative to unmixed condition. Red indicates upregulation and blue indicates downregulation. Genes with significantly different (*P* < 0.05, two-sided t-test) expression levels between the two conditions are indicated by asterisks. Bar charts comparing the Gibbs free energy changes of hydrogen- and formate-mediated propionate oxidation under unmixed and mixed conditions at day 2 (A) and day 8 (B). Mixing makes the hydrogen-mediated pathway more thermodynamically favourable at both time points, with smaller changes for formate. Heatmaps below show changes in hydrogenase and formate dehydrogenase gene expression, with red indicating upregulation and blue indicating downregulation.

Lower propionate consumption at 250 rpm suggested that the cocultures were at earlier growth stages than those at 0 and 130 rpm on both days 2 and 8, complicating comparisons across mixing conditions. Nevertheless, relative to 0 rpm, we still observed more negative ΔG values for both H₂- and formate-mediated pathways, accompanied by increased expression of hydrogenases and formate dehydrogenases (Figure S4A and S4B; Table S6). On day 8, the decrease in ΔG was greater for the H₂-mediated pathway (8.14 kJ/mol) than for the formate-mediated pathway (4.87 kJ/mol). These thermodynamic and transcriptional differences should therefore be interpreted cautiously, as they may reflect both mixing effects and associated differences in physiological stage.

Based on the comparison between 0 and 130 rpm cocultures, which were at similar growth stage on both days 2 and 8 based on propionate consumption, H₂-mediated propionate oxidation showed a greater change in thermodynamic favourability than the formate-mediated pathway. This difference likely reflects stronger transport limitations for H₂ near the cell surface: its surface-to-bulk concentration ratio ranged from 1.28 to 2.16, compared with only 1.00 to 1.03 for formate (Figure S5; Table S7). A similar trend was observed at 250 rpm, although differences in growth stage limit direct comparison with the other treatments. The thermodynamic predictions were generally consistent with the transcriptional responses, except for the weakly expressed Hyb2 and Fdn2 paralogs, which often exhibited expression patterns distinct from those of the dominant Hyb1 and Fdn1 complexes.

### Sensitivity of model predictions to transport and biological parameters

To evaluate model robustness and sensitivity to transport and biological parameters, a one-at-a-time sensitivity analysis was performed by independently varying uncertain parameters relative to a baseline scenario parameterized using experimental measurements. Across all tested parameters and scenarios, the calculated ΔG of the H₂-mediated pathway was consistently more sensitive to parameter uncertainty than that of the formate-mediated pathway (Figure 4; Table S8). The greatest conceptual sensitivity concerned the estimation of the liquid-side mass-transfer coefficient (*k_L_*). In the baseline scenario, *k_L_* was calculated using the stir-bar tip velocity as the characteristic fluid velocity, implicitly assuming that suspended cells experienced the same relative fluid velocity. To represent the extreme case of zero cell–liquid slip, in which cells move with the surrounding liquid and experience no relative advection, we also evaluated a diffusion-only scenario. Removing the advective contribution had the greatest effect at 130 rpm, shifting the calculated ΔG of the H₂-mediated pathway toward less favourable values by 2.47 kJ/mol on day 2 and 1.66 kJ/mol on day 8. The resulting ΔG values were −1.14 and −2.12 kJ/mol, respectively. Under this scenario, the H₂-mediated pathway was predicted to be more thermodynamically favourable under the unmixed condition than at 130 rpm, contrary to the observed transcriptomic response. These results therefore support the inclusion of advection in the mass-transfer model.

**Figure 4.**
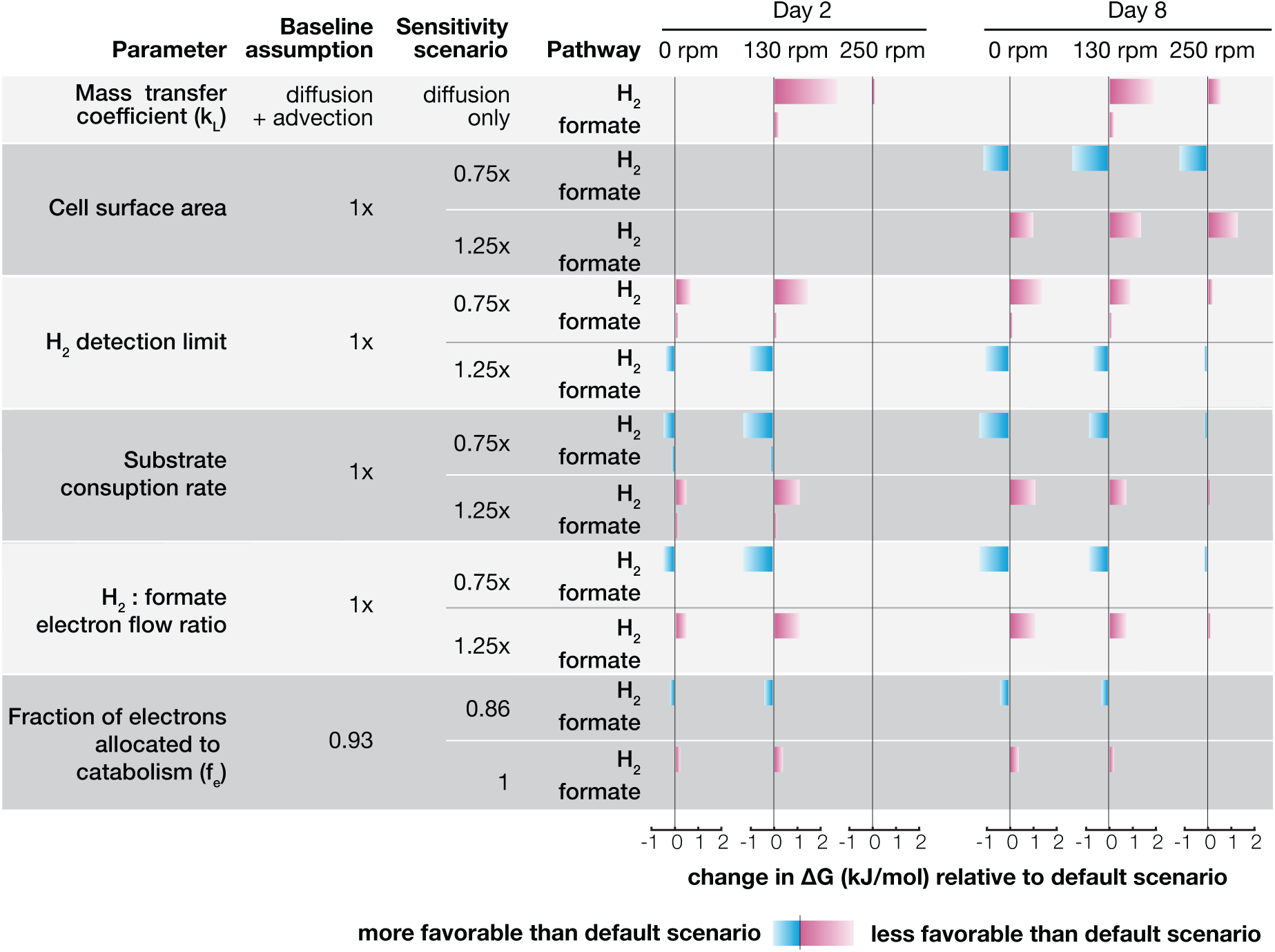
One-at-a-time sensitivity analysis of model-derived Gibbs free-energy changes for H₂- and formate-mediated propionate oxidation. Bars show the change in calculated ΔG relative to the baseline scenario parameterized using experimental measurements for day 2 and day 8 under 0, 130, and 250 rpm. Magenta and blue bars indicate positive (less favourable) and negative (more favourable) deviations from the baseline scenario, respectively. Separate rows are shown for the H₂- and formate-mediated pathways. Sensitivity analysis showing how changes in six model parameters affect the calculated Gibbs free energy of H_2_- and formate-mediated propionate oxidation across mixing conditions on days 2 and 8. Horizontal bars indicate changes relative to the baseline scenario, with magenta representing less favourable and blue representing more favourable conditions. The H_2_-mediated pathway generally shows greater sensitivity than the formate-mediated pathway.

The largest measurement uncertainty arose from the assigned H₂ concentration when H₂ was not detected on GC at day 8. Varying the assigned detection-limit concentration by ±25% changed the calculated ΔG of the H₂-mediated pathway by approximately 1 kJ/mol but had no effect on the formate-mediated pathway. Uncertainty of the remaining transport and biological parameters altered the ΔG of the H₂-mediated pathway by up to 1.5 kJ/mol, with consistently smaller effects on the formate-mediated pathway. This greater sensitivity reflects the stronger dependence of H₂ surface concentrations on transport and biological conditions. Despite these quantitative changes, the principal conclusions remained unchanged: across nearly all tested scenarios, mixing shifted the calculated ΔG of the H₂-mediated pathway toward more favourable values relative to the corresponding unmixed condition.

### The relative use of H_2_- and formate-mediated electron transfer in full-scale anaerobic digesters

To assess whether the relative use of H₂- and formate-mediated interspecies electron transfer also vary in real-world ecosystems, we analysed the expression of formate dehydrogenases and hydrogenases in individual syntrophs using published time-series metatranscriptomic datasets from ten full-scale anaerobic digesters ^54,55^. All digesters were continuously operated, suspended-growth systems, with multiple metatranscriptomes collected from each digester at different time points. We calculated formate dehydrogenase-to-hydrogenase transcript abundance ratios, as a proxy of the relative use of H₂- and formate-mediated interspecies electron transfer for each of the four syntrophs: *Pelotomaculum schinkii*, *Syntrophobacterium fumaroxidans*, *Syntrophus aciditrophicus*, and *Syntrophomonas wolfei* (Table S9-S10). The transcript ratios varied over time for each syntroph within individual digesters (Figure 5), suggesting that the relative use of the two electron-transfer pathways changes even within the same system. For example, in the digester with 14 time-series metatranscriptomes, the ratios ranged from 2.6 to 9.2 for *P. schinkii*, 2.8 to 8.1 for *S. fumaroxidans*, 3.0 to 13.0 for *S. aciditrophicus*, and 0.8 to 3.7 for *S. wolfei*. Redundancy analysis using operational data of this digester (Table S11) indicated that higher formate dehydrogenase-to-hydrogenase transcript ratios were associated with longer hydraulic retention times and higher concentrations of volatile fatty acids and nitrite, whereas soluble chemical oxygen demand, ammonia, and nitrate showed opposite associations (Figure S6). The ratios also differed across digesters, likely reflecting the combined influence of multiple environmental and operational factors on syntrophic metabolism. Ratios exceeded 1 for most syntrophs in most digesters but fell below 1 in some samples, suggesting that formate-mediated electron transfer is preferred over H_2_ under most conditions. Further investigation is needed to identify the environmental and operational factors governing this variation.

**Figure 5.**
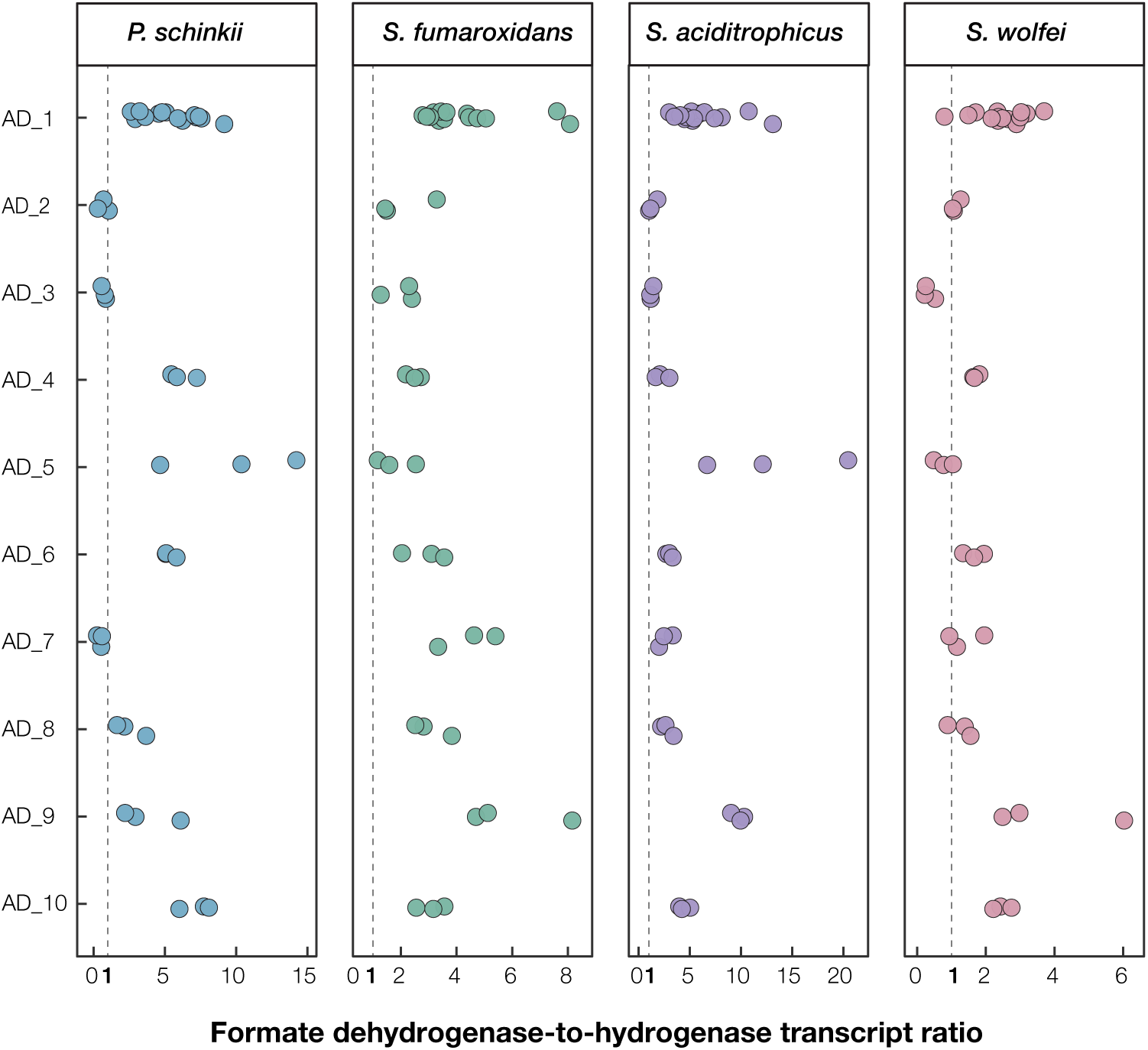
Formate dehydrogenase-to-hydrogenase transcript ratios in four syntrophs across ten full-scale anaerobic digesters. Each point represents a syntroph-specific transcript ratio from one metatranscriptome. Dashed vertical lines indicate a transcript ratio of 1. Points are vertically jittered to reduce overlap. Four-panel scatter plot showing formate dehydrogenase-to-hydrogenase transcript ratios for four syntrophic species across ten anaerobic digesters. Each dot represents one metatranscriptomic sample, with dashed vertical lines marking a ratio of 1. The distribution of ratios varies among species, digesters, and samples within the same digester.

## Discussion

### Fluid flow reshapes the local thermodynamic environment of interspecies electron transfer

This study shows that fluid flow influences interspecies electron transfer during syntrophic propionate oxidation by modulating H₂ and formate concentrations near *Pelotomaculum* cell surface and, consequently, the cell-surface ΔG. Although the energetic differences induced by mixing were only several kilojoules per mole of propionate, shifts of this magnitude are physiologically meaningful because syntrophic propionate oxidation often yields less than 10 kJ/mol propionate ^60^. More broadly, many syntrophic microorganisms operate close to the thermodynamic limit ^2,5,61^. Small changes in electron-carrier concentrations can therefore shift the energetic advantage of alternative electron-transfer pathways.

The effects of fluid flow can be interpreted using the relationship 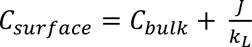. First, fluid flow influences external mass transfer, represented by the liquid-side mass-transfer coefficient, *k_L_*. Experimental studies with suspended microorganisms have shown that fluid flow enhances the transfer of compounds spanning a wide range of molecular diffusivities ^22,23^, while modelling studies suggest that this enhancement results from continuous renewal of the liquid adjacent to the cell ^62^. Accordingly, for a given *J* and *C_bulk_*, mixing-induced increases in *k_L_* promote H₂ and formate removal from the syntrophic cell surface, lowering *C*_surface_ and reducing the thermodynamic burden on propionate oxidation. Second, fluid flow can alter the production fluxes (*J*) of H₂ and formate, by affecting the soluble-substrate uptake rate ^22^ as well as the initiation and propionate degradation rate ^26^. Third, fluid flow can modify *C_bulk_*by regulating transport through the liquid phase, gas–liquid exchange of H₂, and the spatial separation between syntrophs and methanogens, which strongly influences syntrophic interactions and metabolite exchange ^24,63^. Thus, the energetic response to fluid flow therefore reflects the combined effects of near-cell transport, H_2_ and formate production by *P. schinkii*, and their consumption by *M. hungatei*.

The predicted thermodynamic shifts were broadly consistent with changes in gene expression. At 130 rpm, H₂-mediated propionate oxidation became more favourable, whereas the formate-mediated pathway changed little or became slightly less favourable. This shift was accompanied by upregulation of the major hydrogenases and downregulation of formate dehydrogenases. Although differences in growth stage complicate comparison with the other mixing conditions, at 250 rpm, both pathways became more favourable, consistent with increased expression of both enzyme groups. Together, these findings suggest that *P. schinkii* adjusts its electron-transfer machinery in response to flow-induced changes in the local thermodynamic environment. However, greater thermodynamic favourability does not necessarily translate into higher metabolic flux. Despite more favourable calculated ΔG values for both pathways at 250 rpm, the overall propionate oxidation rate was approximately halved. Strong mixing may also affect cellular physiology and growth ^26^. Thus, the responses observed at 250 rpm may reflect effects of mixing beyond changes in electron-transfer thermodynamics.

### H₂-mediated electron transfer was more sensitive to hydrodynamic conditions than formate

We found that the energetics of H₂-mediated propionate oxidation responded more strongly to mixing than those of formate-mediated oxidation, primarily because the dissolved H₂ pool is much smaller. The relative differences between surface and bulk concentrations can be described as 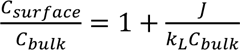. Although H₂ and formate differed by only threefold in production flux (*J*; e.g., 1.02 × 10⁻¹² versus 3.11 × 10⁻¹² µmol/(s·cell) at 0 rpm) and mass-transfer coefficient (*k_L_*; e.g., 0.015 m/s versus 0.005 m/s at 0 rpm) across the examined conditions (Table S2), bulk dissolved H₂ concentration (*C*_bulk,H2_, ∼0.1 µM) was approximately 100–200 times lower than formate concentration (*C*_bulk,formate_, ∼10 µM), consistent with previous modelling results ^31^. Comparable surface accumulation caused a much larger relative increase in H_2_ concentration because of the much smaller denominator (*C*_bulk,H2_). This difference was reflected in the modelled surface-to-bulk concentration ratios, which ranged from 1.011 to 2.407 for H₂ but only from 1.001 to 1.031 for formate (Table S7). Thus, bulk formate closely approximated the concentration at the producing-cell surface, whereas bulk H₂ substantially underestimated the local surface concentration. This agrees with previous studies showing that the low H₂ concentrations required for interspecies electron transfer impose stronger external transport constraints than formate ^31,64^.

An additional source of H₂ sensitivity arises from liquid–gas exchange. Unlike formate, H₂ partitions between the aqueous phase and headspace, allowing mixing to influence its gas–liquid mass transfer rate ^65^. In contrast, based on the p*K*_a_ of formic acid (p*K*_a_ =3.75) and the near-neutral pH of the medium, less than 0.1% of total formate was present as volatile, undissociated formic acid; thus, formate remained almost entirely in the aqueous phase. Mixing therefore introduced an additional carrier-specific redistribution process for H₂. Together, the small, dissolved H₂ pool and its exchange with the gas phase make H₂ surface concentrations, and consequently the thermodynamics of H₂-mediated electron transfer, especially sensitive to fluid flow.

### Mass-transfer-constrained thermodynamic modelling links bulk measurements to the local environment experienced by the cell

Microorganisms respond to the environment immediately surrounding the cell rather than to concentrations measured in the bulk liquid. This distinction has been well established for biofilms, granules, flocs, and microbial aggregates ^31,66–69^. However, mass-transfer gradients are not restricted to attached-growth systems and can also generate differences between local and bulk metabolite concentrations in suspended cultures. Previous studies have shown that diffusion, spatial organization, and extracellular transport resistance can decouple local H₂ and formate concentrations from bulk measurements ^31,64,70^. Our results further demonstrate that, even without visible aggregates, H₂ and formate released at the cell surface can generate local concentrations distinct from the bulk. The magnitude of this difference depended strongly on the transported molecule. It was negligible for formate but substantial for H₂. Bulk formate therefore provided a reasonable approximation of syntroph energetics, whereas bulk H₂ could underestimate the cell-surface concentration and make H₂-mediated propionate oxidation appear more energetically favourable than it is locally.

The model nevertheless remains a simplified representation of transport around syntrophs and can be improved in several ways. First, production fluxes of H_2_ and formate were estimated from average propionate consumption over discrete sampling points, although instantaneous rate likely varied. Future experiments with more frequent measurements throughout the growth cycle could enable estimation of Monod kinetic parameters and thereby provide a more accurate description of time-dependent substrate consumption. Second, the baseline scenario assumed that suspended cells experienced the same relative fluid velocity as the surrounding flow, whereas actual slip velocities may be lower. The diffusion-only scenario included in the sensitivity analysis therefore provided a lower-bound estimate of transport in the absence of advection. Despite uncertainty in *k_L_*, H₂ concentration, cell size, and electron partitioning, the main conclusions remained robust: H₂-mediated electron transfer was consistently more sensitive than formate-mediated transfer and mixing generally shifted pathway energetics toward more favourable values. Third, the present model focused on the propionate-oxidizing syntroph *P. schinkii*, because syntrophs operate at the thermodynamic limit and possess higher genetic diversity in electron transfer that allows flexible use of electron-disposal pathways. In contrast, methanogens usually have a larger energetic margin and are less flexible in electron transfer. Future models could explicitly incorporate methanogen-side transport and energetics, together with gas–liquid mass transfer, to provide a more complete description of syntrophic electron transfer.

### Ecological implications

Environmental microbiology has traditionally focused on the effects of temperature, pH, substrate availability, and redox conditions on microbial activity. In comparison, hydrodynamic conditions remain relatively understudied, despite their potential to regulate metabolite transport, microscale concentration gradients, and spatial interactions ^71–74^. These processes are especially important for syntrophy, whose feasibility depends on rapid removal of H_2_ and formate. Previous studies have shown that H₂- and formate-mediated interspecies electron transfer depends on both cell organization and the surrounding transport environment ^24,70,75,76^. Our results extend these observations by demonstrating that fluid flow influences not only the overall rate of syntrophic metabolism but also the relative thermodynamic favourability and transcriptional regulation of alternative electron-transfer pathways. Hydrodynamic conditions may therefore shape microbial metabolic strategies and should be considered an important ecological factor.

These findings also have implications for engineered systems such as anaerobic digesters. Although mixing is primarily intended to distribute substrates, biomass, and heat, greater mixing intensity does not consistently improve methane production or process stability ^77–80^. Excessive shear can disrupt microbial aggregates and impair syntrophic associations ^81,82^. Our results provide an additional mechanistic perspective by suggesting that fluid flow may alter the microscale transport and thermodynamic constraints governing syntrophic electron transfer. Mixing strategies should therefore balance efficient bulk transport, biomass retention, and favourable microscale conditions for microbial interactions. Because rotational speed tested in serum bottles is not directly comparable across reactor scales, the mixing conditions used here should be interpreted as distinct laboratory hydrodynamic regimes rather than direct analogues of full-scale operation. Pilot- and full-scale studies are needed to translate these mechanistic insights into practical mixing strategies.

In conclusion, this study shows that fluid flow can reshape syntrophic metabolism by altering the near-cell electron-carrier concentrations and pathway-specific energetics. H₂-mediated propionate oxidation was more sensitive to hydrodynamic conditions than formate-mediated pathway. The corresponding transcriptional responses indicate that *P. schinkii* flexibly adjusts its electron-transfer machinery in response to mixing-induced changes in the thermodynamic favourability of H_2_- and formate-mediated electron transfer. Future work should move beyond the estimates of transport around *P. schinkii* by directly resolving cell-scale fluid velocities and methanogenic uptake kinetics. Overall, integrating these improvements will better predict how hydrodynamic conditions regulate syntrophic interactions in natural environments and engineered anaerobic systems.

## Acknowledgement

The staff at the Roy J. Carver Biotechnology Center at the University of Illinois Urbana-Champaign are gratefully acknowledged for the high throughput sequencing assists and formate measurements. We are grateful to Dr. Yoichi Kamagata and Mr. Chao-Jui Chang for their feedback.

## Data availability

Raw transcriptomic data from this study are available in the NCBI Sequence Read Archive under accession PRJNA1479792.

