## Supplementary Information for "A mass-transfer-constrained thermodynamic model reveals how fluid flow reshapes interspecies electron transfer during syntrophic propionate oxidation"

Supplementary Figures

Supplementary Methods

### Supplementary Figures

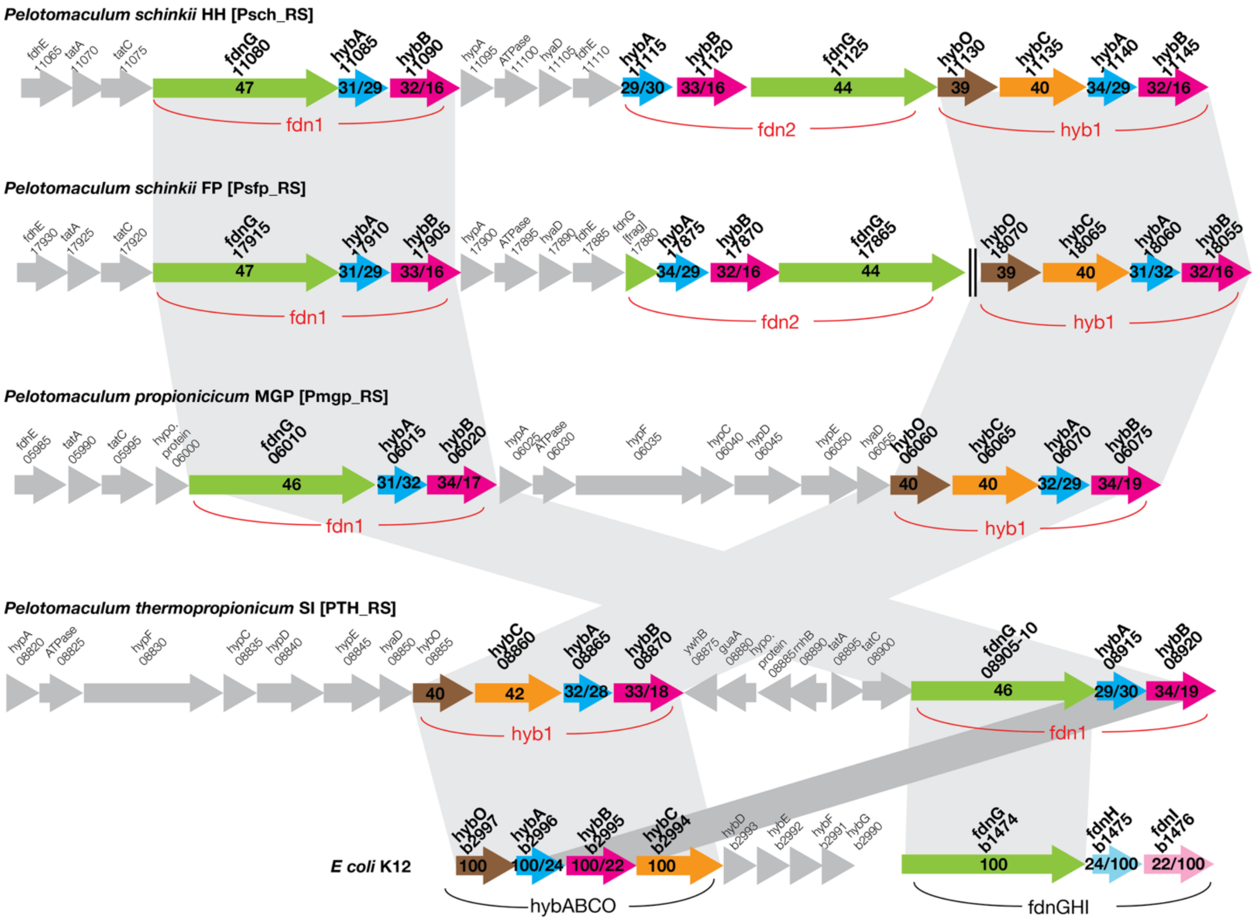


**Figure S1. Conservation and sequence relationships of hyb–fdn gene neighborhoods in propionate-degrading Pelotomaculum species.** Gene neighborhoods containing FdnG-like formate dehydrogenase modules and HybABCO [NiFe]-hydrogenase modules are shown for Pelotomaculum schinkii HH, P. schinkii FP, P. propionicicum MGP, and P. thermopropionicum SI, with the corresponding Escherichia coli fdnGHI and hybABCO loci included as references. The P. schinkii FP locus is encoded on the minus strand. The double vertical line indicates a contig boundary; the flanking genes occur at the end and beginning of separate contigs, respectively. Numbers within FdnG-like proteins indicate percent amino-acid identity to E. coli FdnG. Paired values within Fe–S proteins indicate identity to E. coli HybA/FdnH, and paired values within membrane proteins indicate identity to E. coli HybB/FdnI. Gray arrows indicate neighboring or accessory genes. Gene lengths are drawn approximately to scale.

**Alt-text:** Gene neighborhood diagrams comparing formate dehydrogenase and hydrogenase gene clusters in four *Pelotomaculum* strains and *E. coli*. Colored arrows represent genes investigated in this study, while grey arrows indicate neighboring genes. Numbers within colored arrows show amino acid sequence identities.

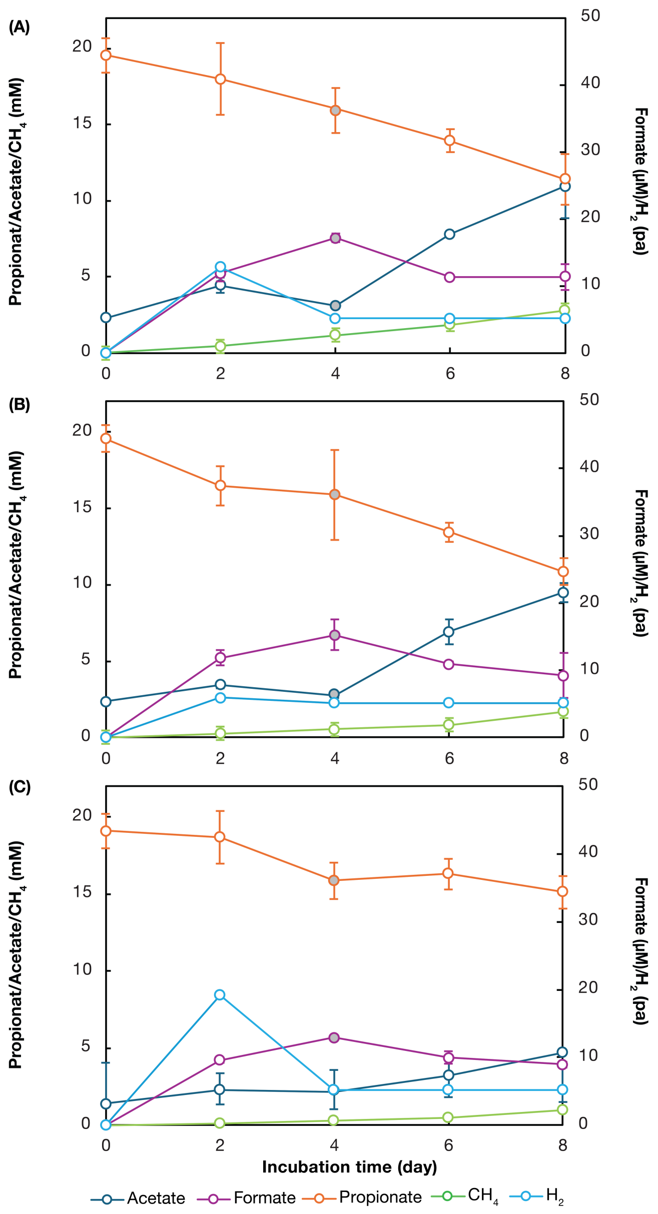


**Figure S2. Metabolite dynamics during syntrophic propionate oxidation at (A) 0, (B) 130, and (C) 250 rpm.** Propionate, acetate, and $CH_4$ concentrations are plotted on the left y-axis, while formate and $H_2$ concentrations are plotted on the right y-axis. Liquid samples from day 4 were accidentally lost during the main experiment; therefore, an additional incubation was conducted under identical conditions to obtain propionate, acetate, and formate measurements for this time point (grey filled circles). Data are presented as means ± SD from three independent cultures (N = 3), except for formate measurements (N = 2).

**Alt-text:** Three-panel line graphs showing metabolite concentrations over eight days at 0 rpm (A), 130 rpm (B), and 250 rpm (C). Propionate decreases while acetate and methane increase, with slower changes at 250 rpm. Hydrogen and formate accumulate early and subsequently decrease.

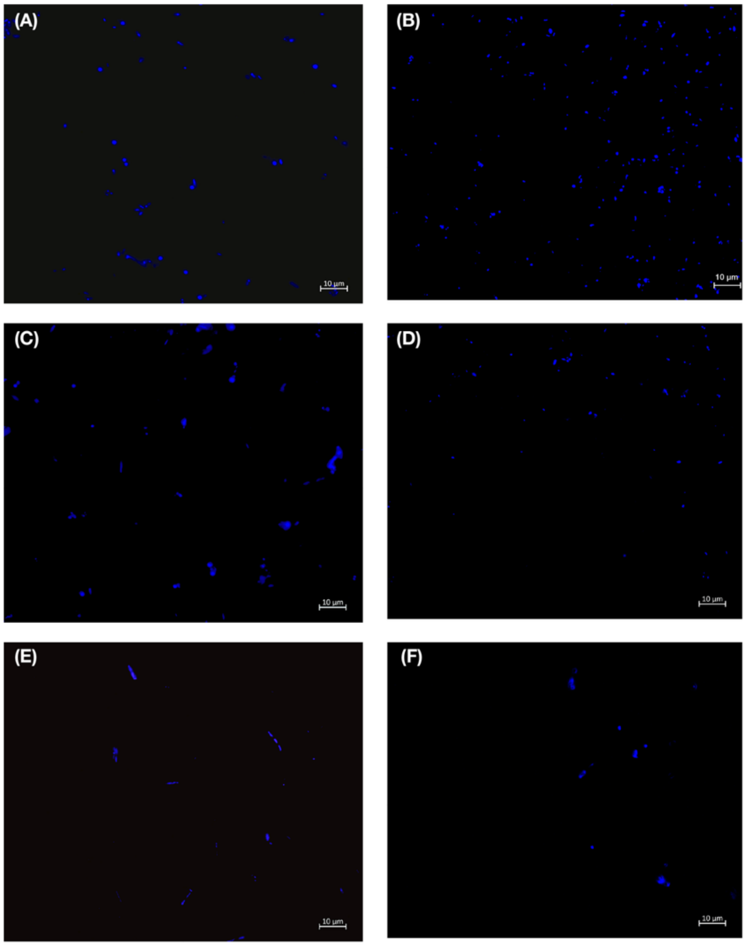


**Figure S3. DAPI-stained cocultures of Pelotomaculum schinkii and Methanospirillum hungatei under different mixing conditions and growth stages.** Representative fluorescence micrographs are shown for cultures incubated without mixing at (A) day 2 and (B) day 8, at 130 rpm at (C) day 2 and (D) day 8, and at 250 rpm at (E) day 2 and (F) day 8.  Scale bars, 10 µm.

**Alt-text:** Six fluorescence micrographs showing DAPI-stained cocultures of *P. schinkii* and *M. hungatei* at 0, 130, and 250 rpm on days 2 and 8. Blue fluorescent cells are dispersed across dark backgrounds, with no visible cell aggregates under any condition.

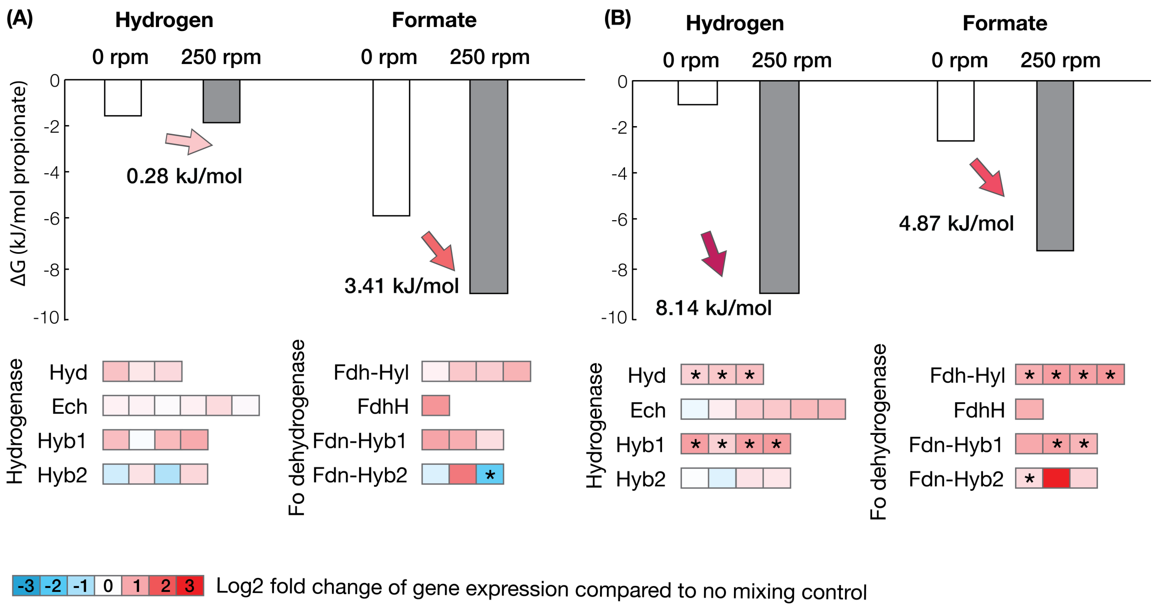


**Figure S4. Mixing-dependent changes in the thermodynamic favorability and transcription of $H_2$- and formate-mediated electron-transfer pathways in *P.* schinkii at 250 rpm.** Bars show the calculated Gibbs free-energy change of propionate oxidation through $H_2$- and formate-mediated pathways under unmixed condition (white) and mixed at 250 rpm (dark grey). Panels compare day 2 (A) versus day 8 (B). Arrows indicate the change in ΔG caused by mixing relative to unmixed control, calculated as $\Delta G_{unmixed}$ - $\Delta G_{mixed}$ ; positive values indicate that mixing made ΔG more negative and thus more favourable, whereas negative values indicate reduced favourability. Heatmaps below each comparison show the log2 fold change in expression of major hydrogenase and formate dehydrogenase complexes at 250 rpm relative to unmixed condition. Red indicates upregulation and blue indicates downregulation. Genes with significantly different ($P < 0.05$, two-sided t-test) expression levels between the two conditions are indicated by asterisks.

**Alt-text:** Two-panel bar charts comparing Gibbs free energy changes of hydrogen- and formate-mediated propionate oxidation at 0 and 250 rpm on days 2 (A) and 8 (B). Mixing makes both pathways more thermodynamically favourable, particularly on day 8. Heatmaps below show changes in hydrogenase and formate dehydrogenase gene expression, with red indicating upregulation and blue indicating downregulation.

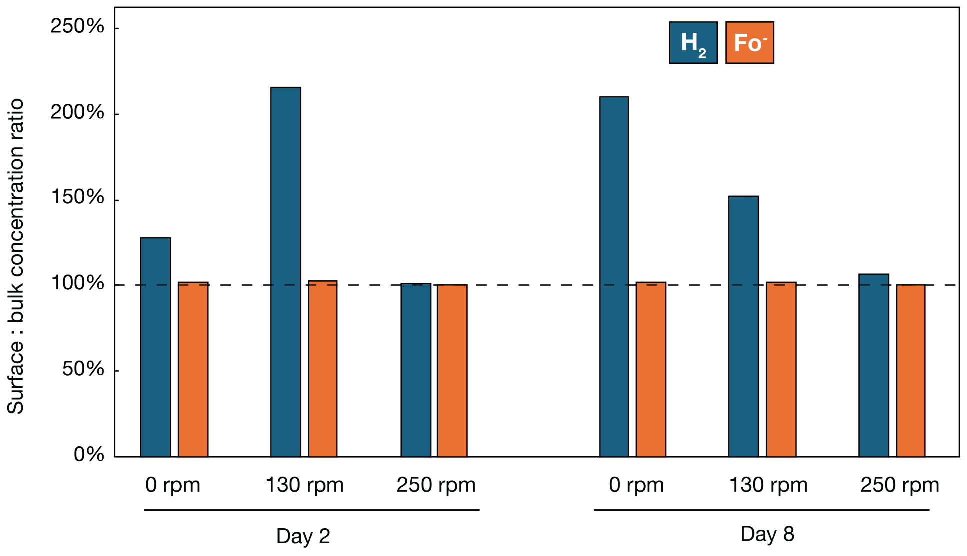


**Figure S5. Surface-to-bulk concentration ratios of $H_2$ and formate under different mixing conditions and growth stages.** The dashed line indicates a surface-to-bulk ratio of 100%.

**Alt-text:** Grouped bar chart comparing surface-to-bulk concentration ratios of hydrogen and formate at 0, 130, and 250 rpm on days 2 and 8. Hydrogen shows greater concentration differences, particularly at 130 rpm on day 2 and 0 rpm on day 8, while formate ratios remain close to 100% across all conditions.

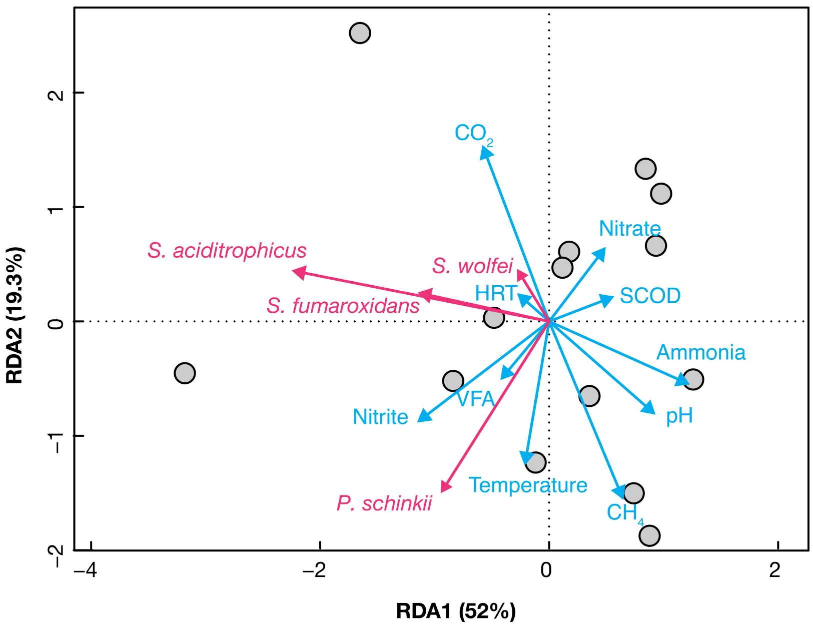


**Figure S6. Redundancy analysis of syntroph-specific formate dehydrogenase-to-hydrogenase transcript ratios and process parameters in an anaerobic digester [54].** Grey circles represent 14 time-series metatranscriptomes; blue arrows indicate process parameters, and pink arrows indicate syntroph-specific transcript ratios. $CH_4$ and $CO_2$ represent biogas percentages. HRT, hydraulic retention time; SCOD, soluble chemical oxygen demand; VFA, volatile fatty acids.

**Alt-text:** Redundancy analysis plot showing relationships between process parameters and formate dehydrogenase-to-hydrogenase transcript ratios for four syntrophic species across 14 samples from an anaerobic digester. Grey circles represent samples, blue arrows represent process parameters, and pink arrows represent species-specific transcript ratios. The first two axes explain 52% and 19.3% of the variation, respectively.

### Supplementary Methods

*Cultivation experiment*

The coculture of *Pelotomaculum schinkii* HH and *Methanospirillum hungatei* JF-1 (DSM 15200) was obtained from the Deutsche Sammlung von Mikroorganismen und Zellkulturen (DSMZ, Braunschweig, Germany). The genomes of the same strain were downloaded from NCBI (accession number GCF_004369205.1 and GCF_000013445.1). Cultures were grown anaerobically in modified Widdel medium prepared following established protocols [1] with slight modifications. The medium contained (per liter): $KH_2PO_4$, 0.15 g; $NH_4Cl$, 0.5 g; $MgCl_2 \cdot 6H_2O$, 0.2 g; $CaCl_2 \cdot 2H_2O$, 0.15 g; $NaHCO_3$, 2.5 g; trace elements solution, 1 ml; Se/W solution, 1 ml; Wolin's vitamin solution, 2 ml; and yeast extract, 0.2 g; resazurin sodium salt, 0.01 g. The trace element solution was based on DSMZ medium 318, except that NaCl was omitted and Se/W was supplied separately at the DSMZ 318 concentration. Propionate was provided at a final concentration of 20 mM. Anaerobic techniques were adapted from previously described methods [2]. All media were prepared under an $N_2/CO_2$ (80:20, v/v) atmosphere, dispensed (45 mL) into 160 mL serum bottles, sealed with butyl rubber stoppers and aluminum crimp caps, and sterilized by autoclaving (121 °C, 20 min). After cooling, media were reduced with $Na_2S$ and cysteine-HCl (0.15 g/L each) from anaerobic stock solutions and supplied with vitamin solution. Cultures were inoculated with 5 mL of actively growing coculture and incubated at 37 °C. Mixing conditions were controlled by magnetic stirring at 0, 130, or 250 rpm using cylindrical stir bars (30 × 6 mm) and a magnetic stirring system. Triplicate bottles were prepared for each condition.

*GC measurement*

Gas samples were analyzed using a GC-2030 gas chromatograph (Shimadzu, Kyoto, Japan) equipped with a 0.5 mL gas sample loop, an SH-Q-BOND PLOT column, an SH-Msieve 5A PLOT column, a thermal conductivity detector (TCD), and a flame ionization detector (FID). Samples first passed through the SH-Q-BOND PLOT column, where $CO_2$ was separated from $H_2$, $O_2$, $N_2$, and $CH_4$. Before $CO_2$ elution, $H_2$, $O_2$, $N_2$, and $CH_4$ were directed to the SH-Msieve 5A PLOT column and detected by TCD; the effluent was subsequently routed to the FID for more sensitive $CH_4$ detection. $CO_2$ eluting later from the SH-Q-BOND PLOT column was directed to the FID channel through a time-programmed valve event. Samples were introduced through a split inlet at 150 °C using helium as the carrier gas. The oven was held at 35 °C for 4 min and then increased to 105 °C at 30 °C/min, giving a total run time of 7 min. The FID was operated at 400 °C with argon make-up gas, $H_2$, and air at 20, 32, and 200 mL/min, respectively. The TCD was operated at 150 °C and 75 mA, with argon as the make-up and reference gas at 15 and 45 mL/min, respectively. Methane and $H_2$ were quantified using external calibration standards. When no discernible $H_2$ peak was detected, $H_2$ concentration was treated as a nondetect and assigned the analyte-specific GC detection limit. According to the Shimadzu GC manual, the detection limit for $H_2$ measured by GC-TCD is 50 ppm, corresponding to approximately 5 Pa, or approximately 2 μM when expressed as a gas-phase molar concentration.

Propionate and acetate in the liquid phase were quantified by GC-FID after acidification and methyl tert-butyl ether (MTBE) extraction. Briefly, 0.2 mL of 0.22 μm-filtered culture supernatant was transferred into a 0.6 mL tube. Iso-butyric acid was used as an internal standard; a 50 mM stock solution was prepared in deionized water and adjusted to approximately pH 2 with phosphoric acid. 4 μL of the internal-standard stock solution were added to each 0.2 mL sample to yield a final internal-standard concentration of approximately 1 mM. Samples were then acidified to pH ≤ 1.5 by adding 5 μL phosphoric acid, followed by addition of 0.1 mL MTBE. The mixture was vortexed for 30 s and allowed to separate into aqueous and organic phases, after which 50 μL of the MTBE phase was transferred to a new vial for GC analysis. Quantification was performed using matrix-matched standards containing the same internal-standard concentration, and peak areas were expressed as analyte/internal-standard ratios. Liquid extracts were analyzed using a GC-2030 liquid-analysis line equipped with an SH-WAX capillary column and an FID. Samples were introduced through a split injector at 200 °C with helium as the carrier gas, a column flow of 5.0 mL/min, and a split ratio of 50. The FID was operated at 400 °C with argon make-up gas, $H_2$, and air at 20, 32, and 250 mL/min, respectively. Volatile fatty acids (VFAs) were quantified against calibration curves generated from matrix-matched standards processed in parallel with samples. Because formate concentrations were near or below the detection limit of our GC method, selected liquid samples were submitted to the Metabolomics team at the Roy J. Carver Biotechnology Center for formate quantification by GC-MS. Samples were diluted 1:1 with 30% $H_3PO_4$ and analyzed using an Agilent 7890B GC coupled to an Agilent 5977A mass selective detector (Agilent Inc, Palo Alto, CA, USA). A 1 μL sample was injected in split mode (15:1) and separated on an HP-INNOWAX column. Helium was used as the carrier gas. The mass spectrometer was operated in positive electron impact mode, and selected ion monitoring targeted m/z 43, 45, 46, 60, and 74. Formate was quantified using an SCFA calibration curve from 0.1 to 10 μM. The limit of quantification and limit of detection for formate were 0.02 and 0.005 μM, respectively.

*RNA sequencing and quality filtering*

Biomass from cocultures was harvested at two time points: day 2 (early phase; ~2 mM propionate consumed) and day 8 (mid-exponential growth phase; ~10 mM propionate consumed) with biological triplicates. Entire cultures were withdrawn from serum bottles using $N_2$-flushed syringes and needles and immediately transferred into 50 mL Falcon tubes pre-flushed with $N_2$ and sealed without delay. Samples were centrifuged at 10,000 × g for 5 min at 4 °C, and the supernatant was decanted, leaving approximately 0.5 mL of residual volume. RNAprotect Bacteria Reagent (QIAGEN, Hilden, Germany) was added (1 mL per sample), and samples were mixed immediately by vertexing for 5 s. After incubation at room temperature (15–25 °C) for 5 min, suspensions were transferred to 1.5 mL microcentrifuge tubes and centrifuged at 5,000 × g for 10 min. The supernatant was removed, and residual liquid was eliminated by briefly blotting the inverted tubes on absorbent paper as required by the RNAprotect reagent. Cell pellets were immediately stored at −80 °C.

RNA extraction was performed after all samples were collected using the RNeasy Mini Kit (QIAGEN, Hilden, Germany), with mechanical lysis using Lysing Matrix E (MP Biomedicals, Irvine, CA, USA) and a MiniBeadBeater-16 (BioSpec Products, Bartlesville, OK, USA; 3450 rpm, 30 s). DNase treatment was performed using the RNase-Free DNase Set (QIAGEN, Hilden, Germany) followed by the TURBO DNA-free Kit (Ambion, Foster City, CA, USA).

Library preparation and sequencing were performed at the Roy J. Carver Biotechnology Center, University of Illinois Urbana–Champaign. Ribosomal RNA was removed using the FastSelect 5S/16S/23S kit (QIAGEN, Hilden, Germany). Strand-specific RNA-seq libraries were generated using the Watchmaker stranded mRNA library preparation kit (Watchmaker Genomics, Boulder, CO, USA). Libraries were pooled, quantified by qPCR, and sequenced on one NovaSeq X Plus lane (10B flow cell) using v1.0 chemistry for 2 × 151 bp paired-end reads.

For the coculture RNA-seq data, raw RNA-seq reads were quality filtered using the standard RNA-seq quality control pipeline developed by the DOE Joint Genome Institute. Briefly, filtering was performed using the BBTools suite (v39.52) [3]. Adapter sequences, low-quality bases, poly-G artifacts, and reads containing ambiguous bases were removed using BBDuk. Optical duplicates were identified and removed using Clumpify. Reads shorter than 51 bp or with an average quality score below 10 were discarded. Reads mapping to host genomes and common contaminants (including human, mouse, and microbial sequences) were removed using BBMap. Ribosomal RNA sequences were filtered using a k-mer–based approach implemented in BBDuk.

*Mass-transfer-constrained thermodynamic modeling*

Based on the classical liquid-film mass-transfer framework [4], we developed a film-theory-based mass-transfer-constrained thermodynamic model to estimate local $H_2$ and formate concentrations at the surface of *Pelotomaculum schinkii* cells and to calculate the corresponding cell-surface Gibbs free energy for syntrophic propionate oxidation. The bulk liquid outside the cell-associated stagnant film was assumed to be homogenous; therefore, measured bulk $H_2$ and formate concentrations were interpreted as the net outcome of syntrophic electron-carrier production, convective-diffusive transport, methanogenic consumption, and, for $H_2$, gas-liquid partitioning. Near the syntrophic cell surface, $H_2$ and/or formate production was assumed to generate concentration gradients across an effective stagnant liquid film of thickness $\delta_i$. The film thickness was related to the liquid-side mass-transfer coefficient by

$$\delta_i = \frac{D_i}{k_{L,i}}$$

, where $D_i$ is the molecular diffusion coefficient and $k_{L,i}$ is the liquid-side mass-transfer coefficient for electron carrier $i$. For each electron carrier $i$, representing $H_2$ or formate, the steady-state flux across the liquid film was described as:

$$J_i = k_{L,i}(C_{surface,i} - C_{bulk,i})$$

where $J_i$ is the cell-surface production flux, and $C_{surface,i}$and $C_{bulk,i}$ are the cell-surface and bulk-liquid concentrations, respectively. Assuming steady state, the diffusive flux $J_i$ across the film equals the $H_2$ or formate production flux $q_i$, allowing the local cell-surface concentration to be estimated as:

$$C_{surface,i} = C_{bulk,i} + \frac{J_i}{k_{L,i}}$$

Bulk formate concentrations were measured in liquid samples by GC-MS. Headspace $H_2$ partial pressure was measured by GC-TCD and converted to dissolved $H_2$ concentration using Henry's law:

$$C_{bulk,H_2} = H_{H_2} p_{H_2}$$

where $H_{\mathrm{H_2}}$ is the Henry's law solubility constant for $H_2$ at 37 °C. In this study, $H_{\mathrm{H_2}}$ was taken as $7.8 \times 10^{-9}$ mol $L^{-1}$ $Pa^{-1}$ [5].

The electron-carrier production flux was estimated from the propionate consumption rate. The propionate consumption rate was calculated from the decrease in propionate concentration between each sampling point and the preceding sampling point two days earlier:

$$r_{pr} = \frac{n_{pr,t-2} - n_{pr,t}}{\Delta t}$$

where $r_{\mathrm{pr}}$ is the total propionate consumption rate. Based on the stoichiometry of syntrophic propionate oxidation, each mole of propionate oxidized releases three electron-carrier equivalents as $H_2$ and/or formate. Therefore, the total production rate of electron carrier $i$was estimated as:

$$R_i = 3r_{\text{pr}} f_e f_i$$

where $f_e$ is the fraction of donor-derived electron equivalents used for catabolism, and $f_i$ is the fraction of electron-carrier flux directed to carrier $i$. The terms $f_{\text{H}_2}$ and $f_{\text{formate}}$ represent the fractions of electron equivalents released as $H_2$ and formate, respectively, with:

$$f_{\text{H}_2} + f_{\text{formate}} = 1$$

The $H_2$/formate partitioning fractions were determined from BES-inhibited incubations, in which methanogenic consumption was suppressed to estimate the relative production of $H_2$ and formate under each mixing condition.

The value of $f_e$ was estimated using a thermodynamic electron-equivalent approach [6]. Electron equivalents released from propionate oxidation were partitioned between energy generation and cell synthesis:

$$f_e + f_s = 1$$

where $f_s$ is the fraction of electron equivalents incorporated into biomass. For each electron-carrier pathway, $f_s$ was calculated as:

$$f_s = \frac{1}{1 + A}$$

where:

$$A = -\frac{\left(\frac{\Delta G_{d-ac}}{\varepsilon} + \frac{\Delta G_{ac-cell}}{\varepsilon}\right)}{\varepsilon \Delta G_e}$$

Here, $\Delta G_{d-ac}$ is the free-energy term associated with converting donor-derived carbon/electron equivalents to the biosynthetic precursor state, $\Delta G_{ac-cell}$ is the free energy required for synthesis of cell material from the precursor state, $\Delta G_e$ is the Gibbs free energy of the catabolic energy-yielding reaction, and $\varepsilon$ is the energy-transfer efficiency. All free-energy terms were expressed on an electron-equivalent basis, in units of kJ per electron equivalent.

The electron fraction used for energy generation was then calculated as:

$$f_e = 1 - f_s$$

For compounds following normal catabolic pathways, the best-fit value for energy transfer efficiency was 0.37 [6]. Using $\varepsilon = 0.37$ , the calculated $f_e$ values were 0.929 for the $H_2$-producing pathway and 0.933 for the formate-producing pathway. Because these values were very similar, a common value of $f_e = 0.93$ was used for subsequent flux calculations. This corresponds to a synthesis fraction of $f_s = 0.07$, consistent with the low biomass yield expected for syntrophic propionate oxidation under methanogenic conditions [7].

The cell-specific production rate of electron carrier $i$ was then calculated as:

$$r_i = \frac{R_i}{N_{\text{syn}}}$$

where $N_{\text{syn}}$ is the number of syntrophic cells, estimated by qPCR. The cell-surface production flux was calculated by dividing the cell-specific production rate by the estimated cell surface area:

$$q_i = \frac{r_i}{A_{\text{cell}}}$$

where $A_{\text{cell}}$ is the surface area of an individual *P. schinkii* cell. Cell surface area was estimated from published electron microscopy-based cell dimensions [8], treating the cells as prolate spheroid.

We also estimated cell-scale mass transfer under two hydrodynamic scenarios in the sensitivity analysis: advection plus diffusion, and diffusion only. The liquid-side mass-transfer coefficient $k_L$ depends on the relative motion between a cell and the surrounding liquid and solute diffusivity. In the present experiments, mixing imposed bulk fluid motion; however, freely suspended cells move with the surrounding liquid and therefore experience substantially less translational cell–liquid slip than indicated by the mixing induced tip velocity. Because the actual relative velocity at the cell surface was not measured, two cell-scale mass-transfer scenarios were evaluated to assess the sensitivity of the calculated surface concentrations and Gibbs free energies to this uncertainty.

For both scenarios, the mass-transfer coefficient $k_L$ for each electron carrier was estimated from the Sherwood number:

$$k_{L,i} = \frac{Sh_i D_i}{2a}$$

where $D_i$ is the diffusion coefficient of $H_2$ ($6 \times 10^{-9}$ m$^2$/s) or formate ($1.9 \times 10^{-9}$ m$^2$/s) in water and $a$ is the volume-equivalent cell radius.

In the diffusion-only scenario, the cell was assumed to move with the surrounding liquid, such that translational cell–liquid slip was negligible. Under this condition, the particle Reynolds number approaches zero and transport around the isolated cell is governed by molecular diffusion. The Sherwood numbers for both hydrogen and formate were therefore set to

$$Sh_i = 2$$

which corresponds to steady diffusion to or from an isolated sphere in a stagnant medium and to the zero-Reynolds-number limit of the Ranz–Marshall correlation [9,10]. The corresponding mass-transfer coefficient was

$$k_{L,i} = \frac{Sh_i D_i}{2a}$$

In the advection-plus-diffusion scenario, the imposed mixing velocity was treated as the characteristic relative velocity between the cell and the surrounding liquid. The Sherwood number was estimated using the Ranz–Marshall correlation ($0 \le Re \le 200$, $0 \le Sc \le 250$) [9,10]:

$$Sh_i = 2 + 0.6 Re^{1/2} Sc_i^{1/3}$$

where $Re$ is the Reynolds number and $Sc_i$ is the Schmidt number:

$$Re = \frac{\rho u (2a)}{\mu}$$

$$Sc_i = \frac{\mu}{\rho D_i}$$

Here, $\rho$ and $\mu$ are the density and dynamic viscosity of water at 37 °C, which are 993 kg/m$^3$ and 0.000691 Pa · s, respectively. The characteristic velocity $u$ was approximated using the stir-bar tip velocity:

$$u = \pi L_{\text{bar}} \left(\frac{\text{rpm}}{60}\right)$$

where $L_{\mathrm{bar}}$ is the stir-bar length (0.03 m). Because stir-bar tip velocity likely exceeds the actual microscale velocity experienced by individual cells, the calculated $k_L$ values should be interpreted as upper-bound estimates. Consequently, the estimated concentration difference between the cell surface and bulk liquid, $J_i/k_{L,i}$, represents a conservative lower-bound estimate of local electron-carrier accumulation.
